# ProBASS – a language model with sequence and structural features for predicting the effect of mutations on binding affinity

**DOI:** 10.1101/2024.06.21.600041

**Authors:** Sagara N.S. Gurusinghe, Yibing Wu, William DeGrado, Julia M. Shifman

## Abstract

Protein-protein interactions (PPIs) govern virtually all cellular processes. Even a single mutation within PPI can significantly influence overall protein functionality and potentially lead to various types of diseases. To date, numerous approaches have emerged for predicting the change in free energy of binding (ΔΔG_bind_) resulting from mutations, yet the majority of these methods lack precision. In recent years, protein language models (PLMs) have been developed and shown powerful predictive capabilities by leveraging both sequence and structural data from protein-protein complexes. Yet, PLMs have not been optimized specifically for predicting ΔΔG_bind_. We developed an approach to predict effects of mutations on PPI binding affinity based on two most advanced protein language models ESM2 and ESM-IF1 that incorporate PPI sequence and structural features, respectively. We used the two models to generate embeddings for each PPI mutant and subsequently fine-tuned our model by training on a large dataset of experimental ΔΔG_bind_ values. Our model, ProBASS (Protein Binding Affinity from Structure and Sequence) achieved a correlation with experimental ΔΔG_bind_ values of 0.83 ± 0.05 for single mutations and 0.69 ± 0.04 for double mutations when model training and testing was done on the same PDB. Moreover, ProBASS exhibited very high correlation (0.81 ± 0.02) between prediction and experiment when training and testing was performed on a dataset containing 2325 single mutations in 132 PPIs. ProBASS surpasses the state-of-the-art methods in correlation with experimental data and could be further trained as more experimental data becomes available. Our results demonstrate that the integration of extensive datasets containing ΔΔG_bind_ values across multiple PPIs to refine the pre-trained PLMs represents a successful approach for achieving a precise and broadly applicable model for ΔΔG_bind_ prediction, greatly facilitating future protein engineering and design studies.

## Introduction

Protein-protein interactions (PPIs) control virtually all crucial processes in the cell including signaling, metabolism, gene expression, cell growth and division, and assembly of large macromolecular complexes^1–3^. Typically, a single protein participates in multiple interactions, contributing to an extensive network of PPIs^4^. Even a single mutation can significantly impact the biding affinity (K_D_) of a PPI, thereby disrupting an existing interaction or creating a new one. Mutations within a single PPI could affect the entire PPI network, leading to alternation in cellular function and often contributing to disease development^5,6^. Mutations in various PPIs have been identified as primary culprits behind conditions like cancer, viral and bacterial infections, and neurodegenerative disorders. Hence, predicting mutational effects on PPI binding affinity helps our general understanding of the disease mechanism and facilitates efforts to design protein-based PPI inhibitors ^6,7^.

The effect of mutation on PPI binding affinity (K_D_) could be determined experimentally by constructing the gene of the mutated protein, expressing and purifying each protein mutant and measuring its affinity to the target protein using one of the established methods such as Isothermal calorimetry (ITC), Surface Plasmon resonance (SPR) or others^8,9^. One such experiment however, requires weeks of work; hence, it is not feasible to measure binding affinity changes for hundreds or more mutations in multiple PPIs. Consequently, computational prediction of changes in free energy of binding (ΔΔG_bind_) presents an appealing alternative to circumvent time-consuming experiments. Over the past decade, many computational tools have emerged for this purpose. These methods are primarily categorized into two groups: sequence-based^10–13^ and structure-based^14–24^ approaches. Both types of methods utilize a set of input features extracted from sequences and/or structures of the interacting partners and train the energy function for ΔΔG_bind_ prediction on these features.

Earlier methods for ΔΔG_bind_ prediction relied only on biophysical atomic-based energy terms such as hydrogen bonding, van der Waals interactions and solvation for ΔΔG_bind_ prediction^19,25,26^. Other methodologies predicted ΔΔG_bind_ utilizing statistical potential energies and coarse-grained protein models ^16,27^. Irrespective of whether these approaches relied on biophysical interactions, statistical potentials or a combination of both, they achieved only a moderate correlation with experimental ΔΔG_bind_ data, exhibiting a Person correlation (R-value) ranging from 0.4-0.6. Incorporation of advanced machine learning techniques into the model building^10,28–31^ improved prediction accuracy achieving correlations of ∼0.7 for single mutations. However, the method accuracy decreased significantly when applied to double and higher-order mutations, producing a correlation of 0.4 or lower ^10^.

Recent years have witnessed a significant breakthrough in the application of artificial intelligence to address diverse biological challenges^32^. Specifically, advanced neural networks such as language models, initially designed for natural language processing, have been adapted to predict a wide array of protein properties ^33–36^. Initially, protein language models (PLMs) were trained on protein sequences and used to predict both global and local protein prediction tasks^33^. Subsequently, PLMs were trained on extensive datasets containing hundreds of thousands of three-dimensional protein structures to forecast the amino acid sequence that ultimately forms this specific protein structure^37^. Recently, a research team from Facebook introduced the ESM-2 model, the most extensive PLM to date, which was trained on 15B protein sequences and stands out for its ability to predict protein structure from sequence data with high accuracy ^38^. The same group also presented an inverse folding model ESM-IF1^39^, which underwent training using a dataset of 12 million protein structures predicted using AlphaFold and has demonstrated exceptional capabilities in reverse engineering protein sequences from their 3D structures^40^. However, these models have not been specifically trained to predict the effect of mutations on binding affinity and are hence less accurate in such predictions compared to other tasks^35^.

Recent studies explored the application of transfer-learning techniques to refine pre-trained PLMs with the aim of enhancing prediction accuracy for specific tasks. For instance, PLMs initially trained on extensive and diverse protein sequence databases have undergone fine-tuning for tasks such as predicting protein secondary structure and the impact of mutations on protein stability ^42,43^ ^41^. This fine-tuning approach has the potential to enhance the accuracy of predictions related to different functional properties of proteins, but it necessitates access to a substantial and consistent dataset to train the model. Until now, researchers have employed the SKEMPI database^42^ as a benchmark to assess the effectiveness of various computational methodologies on ΔΔG_bind_ prediction ^10,14^. This database contains a comprehensive collection of ΔΔG_bind_ values for various PPIs determined through reliable biophysical methods. The limitation of this database is a relatively limited number of data points per one PDB entry and even a smaller number of data points for double mutations, making it sub-optimal for use in deep-learning approaches. Additionally, experimental data in the SKEMPI database was gathered from experiments conducted under various experimental conditions and methodologies and hence is not completely consistent. Our laboratory has amassed an extensive dataset encompassing ΔΔG_bind_ values for tens of thousands of single and double mutations within several serine-protease/inhibitor complexes^43,44^. This dataset was collected using novel methodology that combines yeast surface display technology, deep sequencing, and data normalization on a small set of experimental data collected on purified proteins providing a robust resource for training predictive models for ΔΔG_bind_. Therefore, in the current study, we merged our dataset with the SKEMPI database, resulting in a dataset of nearly 26K experimental ΔΔG_bind_ values, which is substantially larger than previously employed for model training.

To develop a reliable model for ΔΔG_bind_ prediction, in our current work, we combined the sequence-based ESM-2 model and the structure-based ESM-IF1 model into the ProBASS model and retrained it on a large dataset of experimental ΔΔG_bind_ values. The fined-tuned ProBASS model was able to predict ΔΔG_bind_ values with a nearly perfect correlation with experiment when both training and testing were conducted on the same PDB file. This correlation is only slightly reduced when we trained the model on mutational data from multiple PPIs and tested it on a single PPI not included in the training set. Hence, our method proves to be successful in creating a precise and widely applicable model for ΔΔG_bind_ prediction, with the potential for further enhancement as additional experimental data becomes accessible.

## Methods and materials

### Data Preparation

Datasets that were used to train and test the models were derived from the SKEMPI database (including 1868 and 195 single and double mutations, respectively) and our own experimental dataset for 228 single and 13109 double mutations in complex between BPTI and trypsin (PDB ID 3OTJ) and 228 single and 12526 double mutations in complex between BPTI and Chymotrypsin (PDB ID 1CBW)^43^. 44% of our data belonged to the serine-protease/inhibitor complexes while the remaining data belonged to structurally different PPIs. Furthermore, we also tested our model on the deep mutational scanning data reported for the Angiotensin-converting enzyme 2 (ACE2) and Spike protein S1 complex including 358 single mutations in the PPI binding interface area^45^. The full dataset is available as Supporting Information. For each PDB in the dataset, we extracted protein sequences of both chains, protein structures, lists of mutations, and the experimentally measured ΔΔG_bind_ values corresponding to these mutations. If the same complex appeared in both datasets, the data was used only from our own dataset.

### Feature Engineering and machine learning

Two pre-trained language models, ESM-2 and ESM-IF1 were used to extract sequence and structural features, respectively. For each mutation, a sequence for the wild-type PPI and the mutated PPI was extracted and included sequences of both chains. For each mutated and the wild-type sequence, we extracted sequence embeddings using ESM-2 model (1280 embeddings for each position). We then averaged the embeddings over all sequence positions for both the WT and the mutated PPI. Subsequently, we calculated the difference between the embeddings of the mutated complex and those of the wild type complex.

We extracted the structural features from the ESM-IF1 model, Since the structural embeddings describe the positions of the Cα atoms only and hence are very similar for the wild-type and the mutant sequence, we have only used the wild-type embeddings to describe the structural features of the complex. Since the structural embeddings describe the positions of the Cα atoms only and hence are very similar for the wild-type and the mutant sequence, we have only used the wild-type embeddings to describe the structural features of the complex. Thus, 512 structural embeddings were appended to the 1280 sequence embeddings, generating a set of 1792 embeddings for each mutation. Similar to sequence embeddings, structural embeddings were also averaged to represent the whole structure. These structural embeddings were concatenated to the difference of the 1280 sequence embeddings of the wild type and the mutated structural embeddings producing a vector of 1792 features for each mutation in the dataset.

### Training and testing

The Catboost gradient boosting machine learning algorithm ^46^ with RMS as the loss function was used to train our model to predict the effect of mutations on ΔΔG_bind_ values from the embeddings extracted from both sequence and structure of the protein complex. Initially, we focused on protein PPIs with available data for a high number of mutations. We randomly partitioned the mutational data within a single PDB file, allocating 80% for training and 20% for testing. The predicted data was utilized to determine the correlation coefficient between predictions and experimental ΔΔG_bind_ values. We performed a similar training and testing procedure using the whole data set of single mutations containing 132 PDB files, with 80% of the data designated as the training set, and the remaining 20% as a testing. To assess the model’s performance on unseen PDBs, we trained the model on the entire dataset excluding one PDB file and tested the model on the mutations belonging to the unseen PDB. An additional simialr test was done by excluding several PDB files belonging to complexes not homologous to serine protease/inhibitor complexes and testing the model on mutations from these files. To avoid bias for contribution from particular mutations in training, we performed each model training and evaluation several times with different random data allocation to training and testing sets. To determine the minimum training data required for a high correlation between predicted and experimental ΔΔG_bind_ values, we divided the single and double mutation data for the 3OTJ complex into two equal groups: a training set and a test set. Keeping the test set constant, we progressively increased the training set by 5% increments, up to 50%, and then calculated the correlation coefficient for each test. To check the maximum possible correlation between experiment and prediction due to uncertainties in experimental measurements, we generated noise with a mean of zero and a standard deviation based on the experimental error in each experimental measurement. The noise was drawn from a normal distribution and was added to the experimental measurements. RMSE (Root Mean Square error) for each graph was calculated using the equation

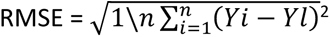

*Yi* :- Experimental determined value for the *ith* data point.

*Yl* :- Predicted value for the *lth* data point.

*n* :- Number of data points.

To compare the results obtained by our model to those of previously developed methods, we utilized three state-of-the-art methods to obtain ΔΔG_bind_ predictions: ESM-IF1^47^, ProteinMPNN^48^, and ThermoMPNN^49^. In ProteinMPNN, we derived the negative log probability score for each mutation and correlated it with the experimentally measured ΔΔG_bind_ values. Similarly, for the model ESM-IF1, we also used the sequence of each PPI and calculated the conditional log-likelihoods for sequences conditioned on a given structure and correlated the results with the experimentally measured ΔΔG_bind_ values. We calculated the change in thermostability (ΔΔG) values for ThermoMPNN by utilizing the wild-type protein complex and the chain identifier. Subsequently, we correlated these results with experimentally measured ΔΔG_bind_ values.

## Results

To build the ProBASS model, we used two state-of-the-start pre-trained PLMs, ESM-2 and ESM-IF1, that contain sequence and structural features, respectively. For each mutation in a particular PPI, we first extracted sequence embeddings from the ESM-2 model, collecting 1280 embeddings per each sequence position (Figure 1). This process was performed for WT and the mutated PPI sequences and the embeddings were subsequently averaged over all sequence positions, effectively condensing the information into 1280 sequence-derived embeddings per mutation. Subsequently, the difference between the embeddings of the mutated complex and those of the WT complex was calculated to reflect the change in features due to mutation. Structural embeddings were obtained for each PPI using the ESM-IF1 model. Since structural embeddings describe positions of the backbone (N, Cα and C atoms) only, which in most cases would be very similar for the WT and the mutant sequence, we only used the WT embeddings to describe the structural features of the complex. Thus, 512 structural embeddings were appended to the 1280 sequence embeddings, generating a set of 1792 embeddings for each mutation (Figure 1). Next, ProBASS was trained using the Catboost algorithm^46^ on various subsets of our large experimental database of ΔΔG_bind_ values that included 2,325 single and 25,840 double mutations in 132 PPIs (see Methods for the details).

**Figure 1.**
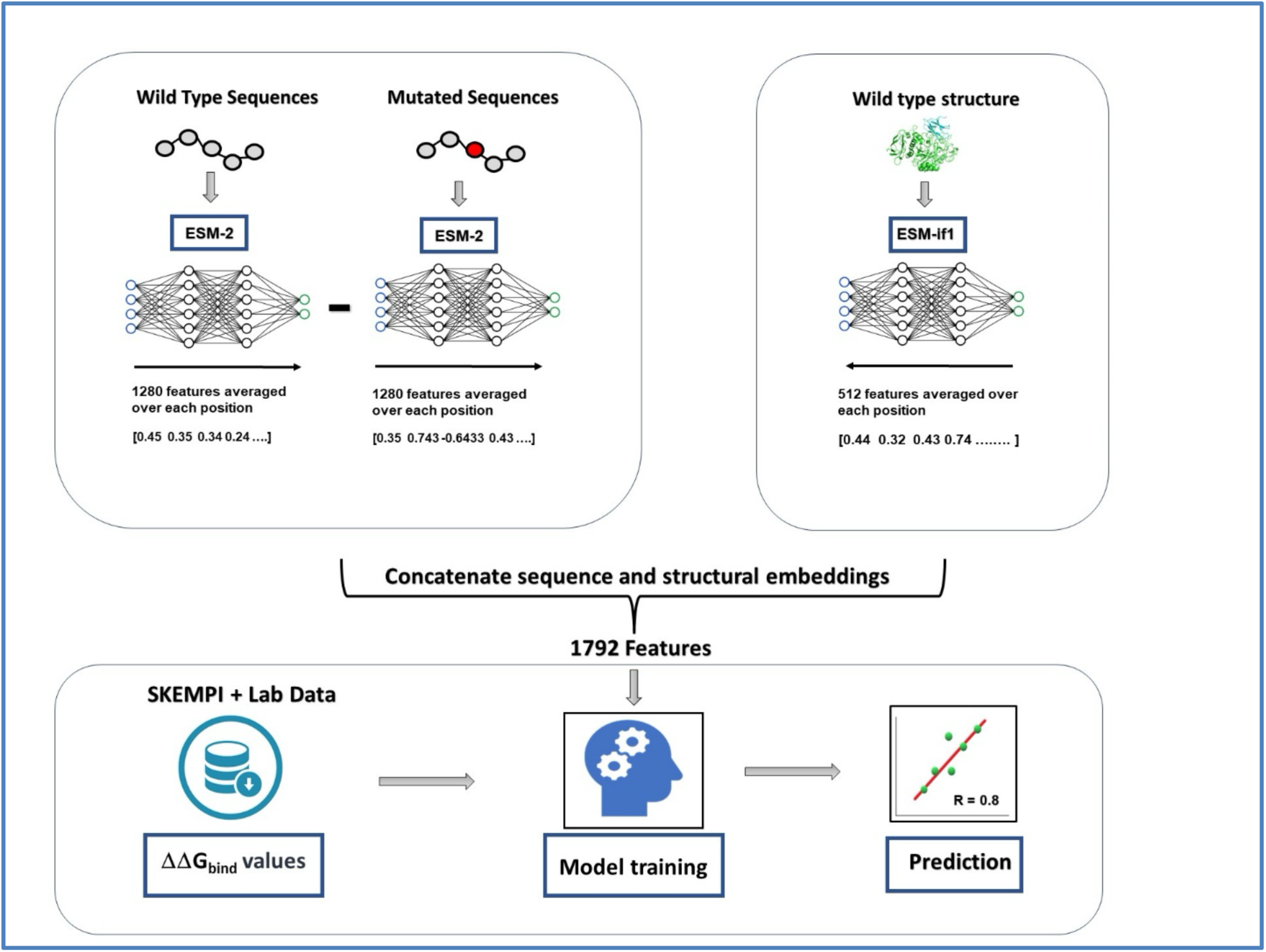
Flow chart for model building. Sequence features were extracted using the EMS2 model and structure features were extracted using the ESM-IF1 model. Embeddings were extracted per residue and later averaged over the sequences of the interacting partners. Both sequence and structure features were concatenated and trained using the Catboost algorithm on the dataset of experimental ΔΔG_bind_ values.

First, we evaluated whether ProBASS could reliably predict ΔΔG_bind_ values on the level of a single PPI. For this purpose, we selected PPIs for which a high number of data points (∼200 mutations) was available since smaller number of mutations would not allow us to perform a reliable model training and evaluation. Data points for one PPI were randomly assigned to the training and the test sets containing 80% and 20% of the data points, respectively. After model training, ΔΔG_bind_ values were predicted for the test set of mutations and the correlation between predictions and experimental values was calculated. To minimize the influence of particular mutations on model training, we repeated the training procedure three times and averaged the correlation coefficient. Our results showed that training and testing on the same PDB produced very high correlations for all tested PDBs ranging from 0.77 – 0.91 with a root mean square error (RMSE) of 1.2 kcal/mol (Figure 2 and Supplementary Figure 1). In comparison, using experimental binding affinity data in colicin/DNAse^50,51^ complexes, we estimated that the maximum possible correlation between experiment and prediction would be ∼0.95 due to uncertainties in each experimental measurement (Supplementary Figure 2). This correlation would be further reduced if experimental binding affinity data were measured by different methods or under different conditions.

**Figure 2.**
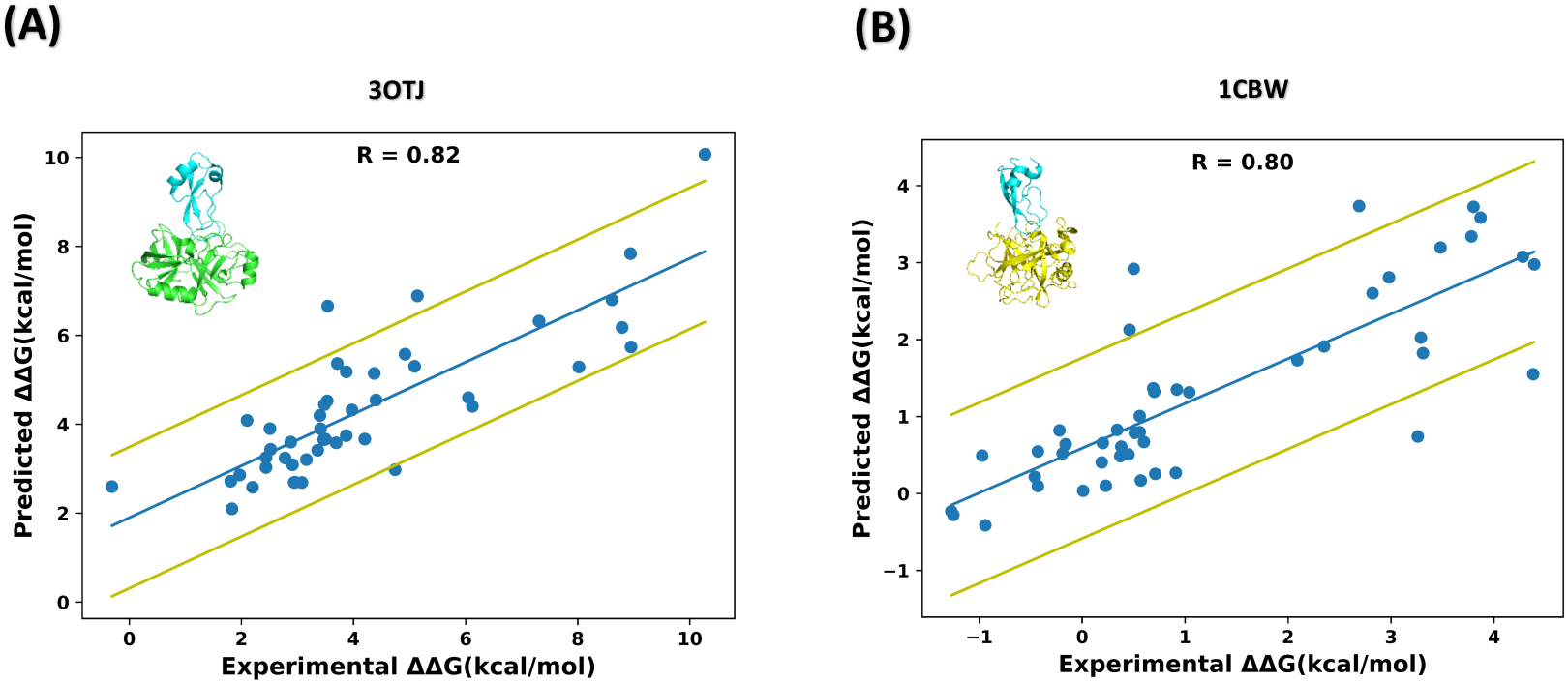
Correlation between experimental and predicted ΔΔG_bind_ values when training and testing on a single PPI: (A) a complex between BPTI and Trypsin (PDB ID 3OTJ) and (B) a complex between BPTI and Chymotrypsin (PDB ID 1CBW). The blue line represents the best linear fit of the data. The yellow lines represent one standard deviation above and below the fitted line.

Next, we tested whether our model could be trained on one PPI and predict ΔΔG_bind_ values in another PPI. For this purpose, we trained the model using the data on the BPTI/bovine trypsin complex (PDB ID 3OTJ) and tested it for predicting ΔΔG_bind_ values for the Proteinase b/ Turkey ovomucoid inhibitor complex (PDB ID 3SGB) (Figure 3A). We observed that in such a case, the correlation between prediction and experiment was reduced considerably to 0.41 with a RMSE of 2.6 kcal/mol, indicating that ΔΔG_bind_ values were heavily dependent on PDB under study and the learning could not be transferred from one complex to another. In attempt to obtain a more generalized model for ΔΔG_bind_ prediction, we decided to perform the training on mutations in multiple PDB files and to test the model on another PDB file not included in the training set. When training was performed on the data for 2135 single mutations from 131 PDB files and testing on 190 mutations belonging to the 3SGB file, the R-value was increased to 0.81 (Figure 3B). Slightly worse correlations were obtained when testing was performed on different PDB files unseen by the model including multiple non-serine protease/inhibitor complexes (Supplementary Figure 3) with the average correlation of 0.68 and RMSE of 1.85 kcal/mol for the six performed tests. These results suggest that training on multiple PDBs could greatly improve the accuracy of predictions on a PDB file not included in training. In addition, we evaluated the ability of our model to reproduce deep mutational scanning data, which measures relative binding affinity when one of the proteins is expressed on the yeast surface. Our model gave a correlation of 0.51 with such semi-quantitative data for the complex between Angiotensin-converting enzyme 2 (ACE2) and the spike protein S1, the complex which shares no homology with any structures in our training dataset (Supplementary Figure 3).

**Figure 3.**
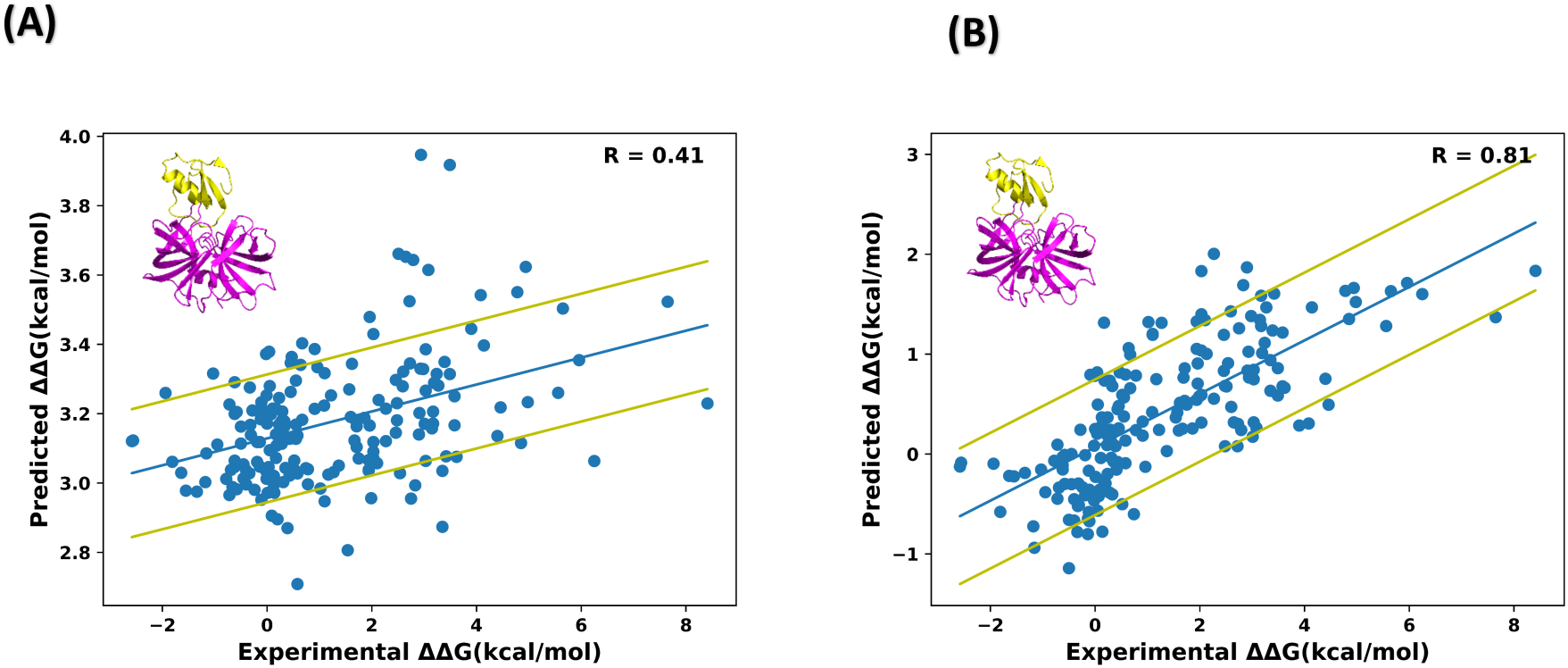
Correlation between predicted and experimental ΔΔG_bind_ values for the Proteinase b/Turkey ovomucoid inhibitor complex (PDB ID 3SGB) for various training scenarios (A) The model was trained on the BPTI/ Trypsin complex (PDB ID 3OTJ). (B) The model was trained on the whole dataset excluding the 3SGB data. The blue line represents the best linear fit to the data. The yellow lines represent one standard deviation above and below the fitted line.

In a further test, we randomly divided our entire single mutational dataset into a training set (80% of mutations) and a testing set (20% of mutations). In such a test, mutations from the same complex could potentially appear in both sets. Subsequently, we examined the correlation between the predicted and experimental ΔΔG_bind_ values in the test set, repeating the procedure five times (Figure 4). Our analysis shows correlation of 0.81 ± 0.02 between prediction and experiment and RMSE of 1.2 kcal/mol, demonstrating high prediction accuracy on the whole dataset of single mutations.

**Figure 4.**
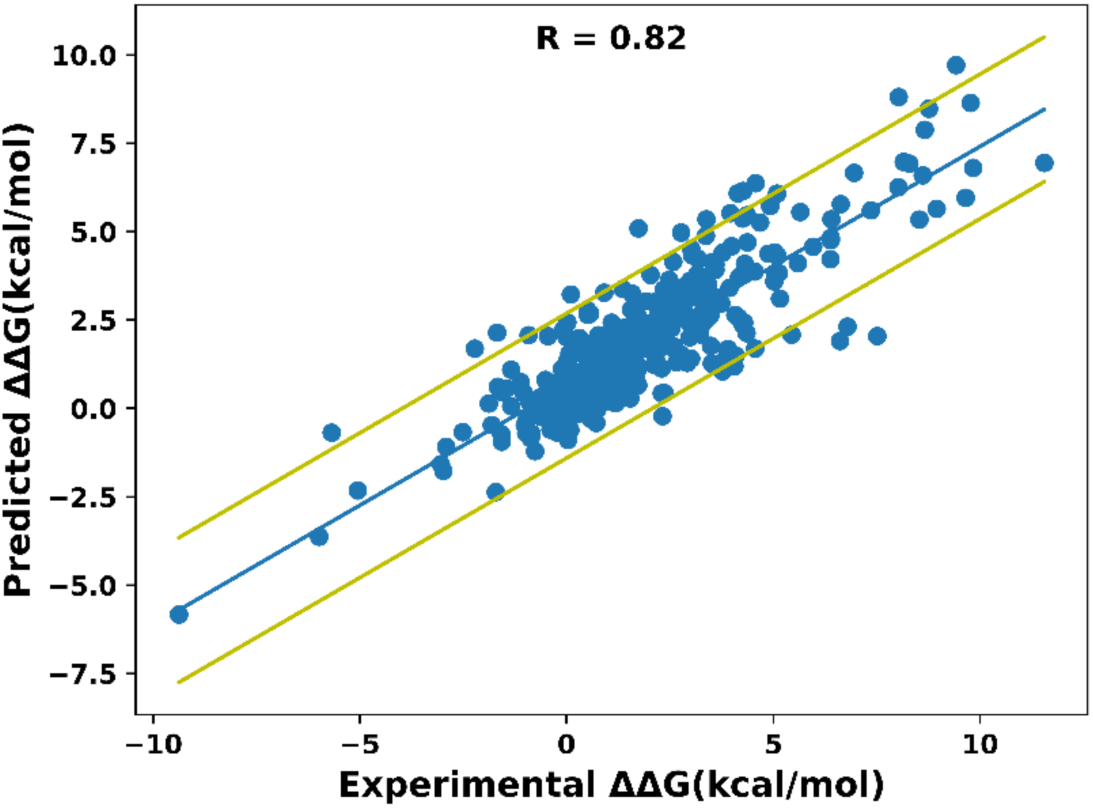
Correlation between experimental and predicted ΔΔG_bind_ values when training and testing was performed on the whole dataset of single mutations. Mutations were randomly allocated into training and testing sets (80% and 20% of data points, respectively), allowing mutations from the same PDB to potentially appear in both sets. The blue line represents the best linear fit of the data. The yellow lines represent one standard deviation above and below the fitted line.

Next, we assessed the performance of our model in predicting ΔΔG_bind_ values for double mutations, a task that is typically considered more challenging in the field of prediction. Again, we initially examined correlation between predicted and experimental ΔΔG_bind_ values when training and testing was performed on the same PPI. Here, we examined two PPIs that contained the data for ∼13,000 mutations each and were able to obtain an R-value of ∼0.7 between prediction and experiment and RMSE of 2.3 kcal/mol (Figure 5A and B). This correlation is slightly lower compared to that obtained for single mutations, yet considerably higher than that reported in previous studies^10^. Just as we examined predictions for single mutations, we trained our model on double mutations in the trypsin/BPTI complex (PDB ID 3OTJ) and tested on the mutations in the chymotrypsin/BPTI complex (PDB ID 1CBW). We obtained a correlation of approximately 0.4 and RMSE of 3.85 kcal/mol between prediction and experiment, similar to the correlation produced in the same protocol for single mutations (Supplementary Figure 5A). Subsequently, following a training approach akin to that used for single mutations, we expanded our model’s training to encompass a wider spectrum of double mutations across different protein complexes. However, we did not see improvement in correlation in this approach very likely due to the limited availability of double mutational data and the dominant impact of the two PDB files with 26,000 mutations (3OTJ and 1CBW) in the training set (Supplementary Figure 5B).

**Figure 5.**
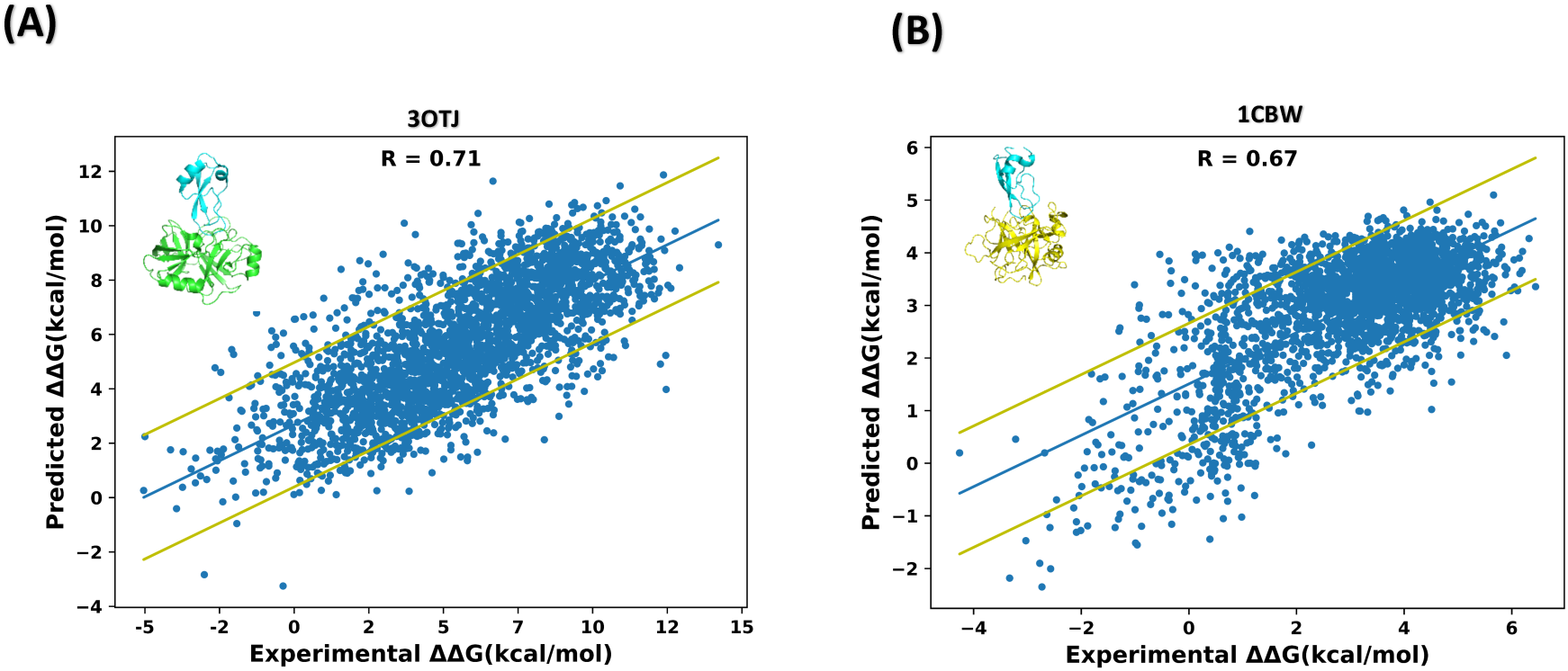
Correlation between experimental and predicted ΔΔG_bind_ values for double mutations (A) training and testing was done on the BPTI/ Trypsin complex (PDB ID 3OTJ). (B) training and testing was done on the BPTI/Chymotrypsin complex (PDB ID 1CBW). The dataset was partitioned randomly with 80% of the data allocated to the training set and 20% to the testing set. The blue line represents the best linear fit to the data. The yellow lines represent one standard deviation above and below the fitted line.

We further evaluated the minimal amount of the training data that would result in high correlation between ΔΔG_bind_ prediction and experiment. For this purpose, we randomly partitioned single and double mutational data within the 3OTJ complex into two equal groups, the training and the test sets. While keeping the testing set constant, we systematically increased the training set in increments of 5% to reach the maximum 50% and computed the correlation coefficient for each protocol. Our results show that correlation is low if the training set is small but rapidly increases and reaches the value of ∼0.7 when approximately 25% of the data is used for training (Supplementary Figure 5).

We next compared the performance of our model with that of other available PLMs such as ESM-IF1^39^ that was trained on billions of protein structures without fine-tuning on ΔΔG_bind_ data and to that of ProteinMPNN^48^, the cutting-edge protein design software that uses graph-based neural network approach to design protein sequences for a particular structure. We additionally evaluated predictions by ThermoMPNN^52^, an expanded version of ProteinMPNN that has been retrained on a wide range of data representing mutational effects on protein stability. Figure 6 shows that ProBASS achieved the highest correlation between the predicted and experimental ΔΔG_bind_ values for both single and double mutations for all tested PDB files with a highest correlation of 0.81 reached for the Proteinase B/ Turkey ovomucoid inhibitor complex (PDB ID 3SGB). Some models performed reasonably well on some PDB files but failed on others. ThermoMPNN which has been retrained to predict the effect of mutations on stability, exhibited the lowest overall accuracy in predicting the effect of mutations on binding. This finding demonstrates that fine-tuning of PLMs for one task does not help in improving predictions for a different task.

**Figure 6.**
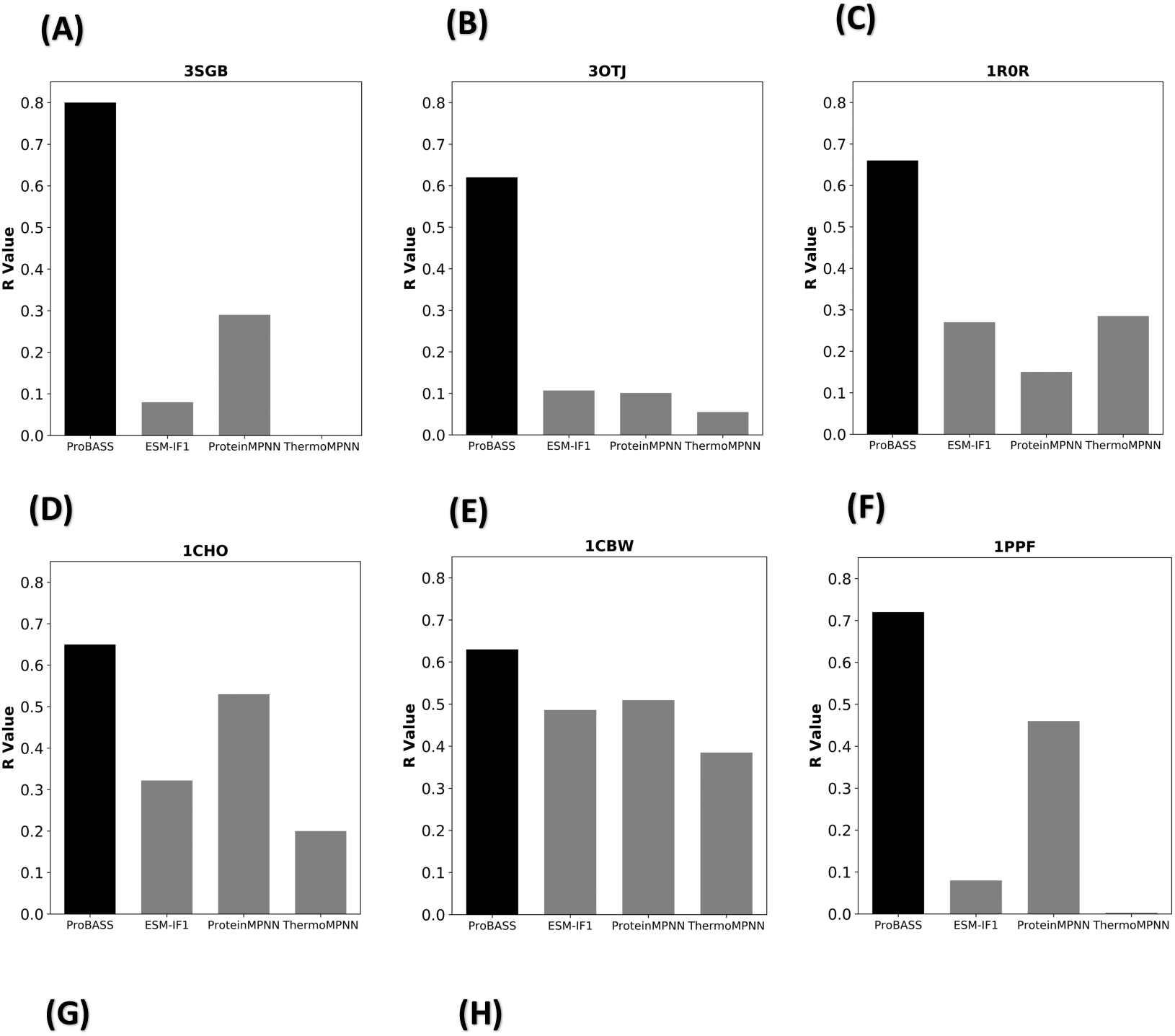
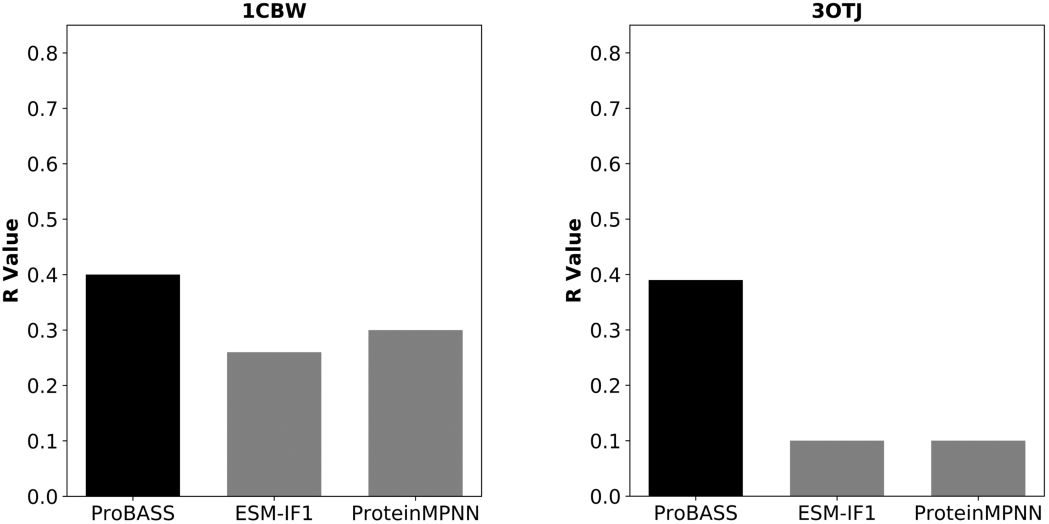
Comparison of performances for our ProBASS model and other available models to predict the effect of mutations on protein binding affinity. Spearman’s correlation coefficients obtained by ProBASS, ESM-IF1, ProteinMPNN and ThermoMPNN on single mutations (A-F) and double mutations (G-H) for different PDB IDs. (A) single mutations for PDB 3SGB. (B) single mutations for PDB 3OTJ (C) single mutations for PDB 1R0R, (D) single mutations for PDB 1CHO (E) single mutations for PDB 1CBW, (F) single mutations for PDB 1PPF. **(G)** double mutations for PDB 3OTJ, (H) double mutations for 1CBW.

## Discussion

In this study we developed a state-of-the art ProBASS model for predicting the impact of mutations on changes in binding affinity by leveraging both sequence and structural data, extracted from the PLMs, ESM2 and ESM-IF1, and fine-tuned this model on a large set of experimental ΔΔG_bind_ data. Our model can predict ΔΔG_bind_ values for single mutations with extremely high accuracy (R-value of 0.8 and higher) when training and testing is done on the same PPI and slightly lower average (R-value ranging from 0.61 to 0.81) when training on multiple PPIs and testing on a different PPI. These results suggest that once a sufficient amount of experimental data has been gathered for training, additional experiments may not be necessary as ΔΔG_bind_ predictions can be made reliably. Although the current dataset contains a high number of mutations belonging to the serine protease/inhibitor complexes (44%) that might bias training toward this PPI structure, we were able to obtain high correlation with experiment also for structurally different PPIs and an excellent correlation if training and testing is performed on the whole dataset containing 132 PDBs. In addition, we proved that a similar strategy could be applied to double mutations if a sufficient amount of mutational data is available, such as in case of PDBs 3OTJ and 1CBW. However, due to the lack of sufficient experimental ΔΔG_bind_ values for double mutations on additional PPIs, it was not possible to generalize our model for double mutations at this point. Moreover, when trained on multiple PDBs but tested on one unseen PDB, correlation was high (R-value of ∼0.7) but the actual predicted and experimental values differed in absolute value (Figure 3B). These results are consistent with the fact that PPI energetic binding landscapes depend on the PPI evolutionary optimality and could differ substantially for even highly homologous complexes^43,44^. Our previous study demonstrated that same mutations could produce highly different effects even in structurally similar PPIs^43^. Thus, when training and testing are done on PPIs with different binding landscapes, low correlation is expected. Training on multiple PPIs however, averages multiple binding landscapes and results in better overall prediction for an unseen PPI. Possible difference in absolute values of ΔΔG_bind_ predictions, however could be explained by the difference in magnitudes of effects in different PPIs as well as different experimental conditions used for collection of the training dataset.

We observe that our approach of using both sequence and structure features and fine-tuning the PLM model for ΔΔG_bind_ prediction produces superior results compared to alternative state-of-the-art methodology based on PLMs as shown on Figure 6. In fact, these three methods achieve very low correlations with experimental data for some of the PDB files (R=0.1 for PDB IDs 3SGB and 1PPF) (Figure 6). This low correlation for some PDB files is likely due to the fact that both ProteinMPNN and ESM-IF1 primarily rely on structural features for training and have not been fine-tuned for ΔΔG_bind_ prediction. In addition, the ESM-IF1 model is trained on individual proteins, potentially limiting its ability to capture the distinctive features of protein complexes. ThermoMPNN that is trained on mutational effects on protein stability exhibited the lowest correlation with experiment, suggesting that fine-tuning on one particular prediction task could only decrease the accuracy of prediction for another task. A few previous studies explored the use of fine-tuning PLMs and Graph neural networks for ΔΔG_bind_ prediction. One such model, ELASPIC2, used two pre-trained neural networks, ProteinSolver^53^ and ProtBert^54^ to generate features and fine-tune them to predict ΔΔG_bind_ among other protein properties. Yet, the reported correlation with experimental data for this model remained low, reaching 0.4 for the SKEMPI dataset. Higher observed correlation in our work could be due to superiority of the ESM-2 and the ESM-If1 models used in current work and a much more comprehensive experimental dataset utilized for fine-tuning.

To understand where further improvements to our model could be implemented, we examined the nature of outlier mutations in six PPIs with the highest number of data points available (Supplementary Figure 6). For each tested PDB file, we first identified the outliers or mutations that lie further than one standard deviation from the best fit. Our analysis shows that outliers depend on the PDB under study and are sometimes but not always conserved in homologous PPIs. We first examined the set of substitutions that were predicted to be more destabilizing than observed experimentally with statistical significance. Among such amino acids were aromatic residues (P and Y) and hydrophobic residues (F, I, L, Y, M, V) that were both significantly enriched as a group among predicted over-destabilizing mutations (binomial P-value < 0.001 and < 10^-5^, respectively). Underrepresented among such mtuations were polar residues (D, E, K, N, Q, R,) with a P-value < 10^-5^. Finally, residues that tend to disrupt secondary structure, Pro, Gly, Asn were depleted in this group (P < 0.001). On the other hand, mutations that were predicted to be more stabilizing than observed experimentally tended to be polar (D, E, K, N, Q, R, P-value < 10^-5^), particularly E and K (P-value< 0.05). Similarly, residues that tend to disrupt secondary structure (P, G, N) were enriched (P-value < 0.001) and hydrophobic residues (F, I, L, F, M, V) were depleted in this population (P-value < 10^-4^). These findings reveal possible bias in the model, which appears to under-reward hydrophobic substitutions relative to polar and secondary structure destabilizing mutants.

Of these, only mutations to proline, were enriched at over 2σ greater than expected. Such mutations are likely to cause local backbone changes that are not reflected in our model. Other frequent outlies include mutations from small to aromatic amino acids, where such mutations are predicted to be over-stabilizing when performed on the surface of the binding interface. Again, such mutations could result in backbone conformational changes and additionally might not be present in sequence alignment used for feature extraction.

In conclusion, in our study, we developed a cutting-edge model ProBASS for predicting the impact of mutations on changes in binding affinity, which leverages both sequence and structural PPI features. Using the combination of ESM2 and ESM-IF1 models for feature extraction proved beneficial, demonstrating their ability to navigate the complex relationship between protein sequences, structure and binding affinity. In addition, fine-tuning the model for prediction of ΔΔG_bind_ values proved crucial in enhancing the predictive power of PLMs, illustrating their ability to adapt to specific tasks. ProBASS could be further improved by retraining on additional experimental data as such data becomes available. This would be especially important for the development of a generalized model for ΔΔG_bind_ prediction for double and higher number of mutations, where experimental data is still scarce. Accurate prediction of ΔΔG_bind_ values by ProBASS enables identification of residues essential for sustaining functional PPIs, understanding the effect of various disease-associated mutations and facilitating a wide range of applications in protein engineering and design.

## Acknowledgements

This study was supported by Israel Science foundation grant 3486/20 (J.M.S.). In addition, J. S. M. acknowledges support from ICRF, NSF/BSF grant 2022685 and the U. of Toronto/HUJI research alliance in protein engineering, and NIH R01CA258274.

## Data and software availability

Software could be downloaded from https://github.com/sagagugit/ProBASS

---

**Supplementary Figure 1:**
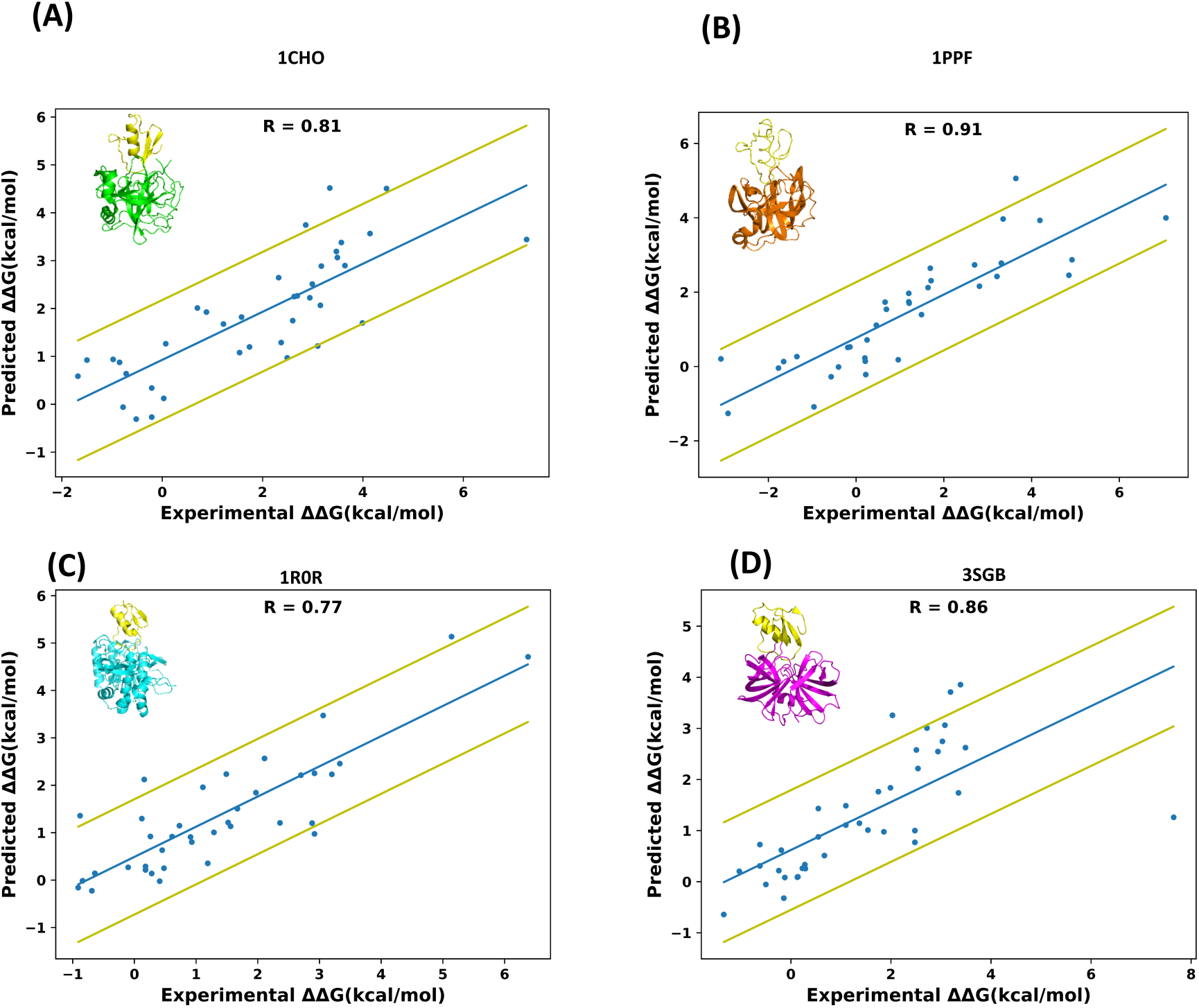
Predicting ΔΔG_bind_ on single PPIs. Correlation between experimental and predicted ΔΔG_bind_ values when training and testing is performed on a single PPI: (A) a complex between Ovomucoid and Alpha-Chymotrypsin (PDB ID 1CHO), (B) a complex between Ovomucoid and Human Leukocyte elastase (PDB ID 1PPF), (C) a complex between Ovomucoid and subtilisin carlsberg (PDB ID 1R0R), (D) a complex between Ovomucoid and subtilisin Proteinase b (PDB ID 3SGB). The blue line represents the best linear fit of the data with the Person correlation R-value given on each graph. The yellow lines represent one standard deviation above and below the fitted line.

**Supplementary Figure 2:**
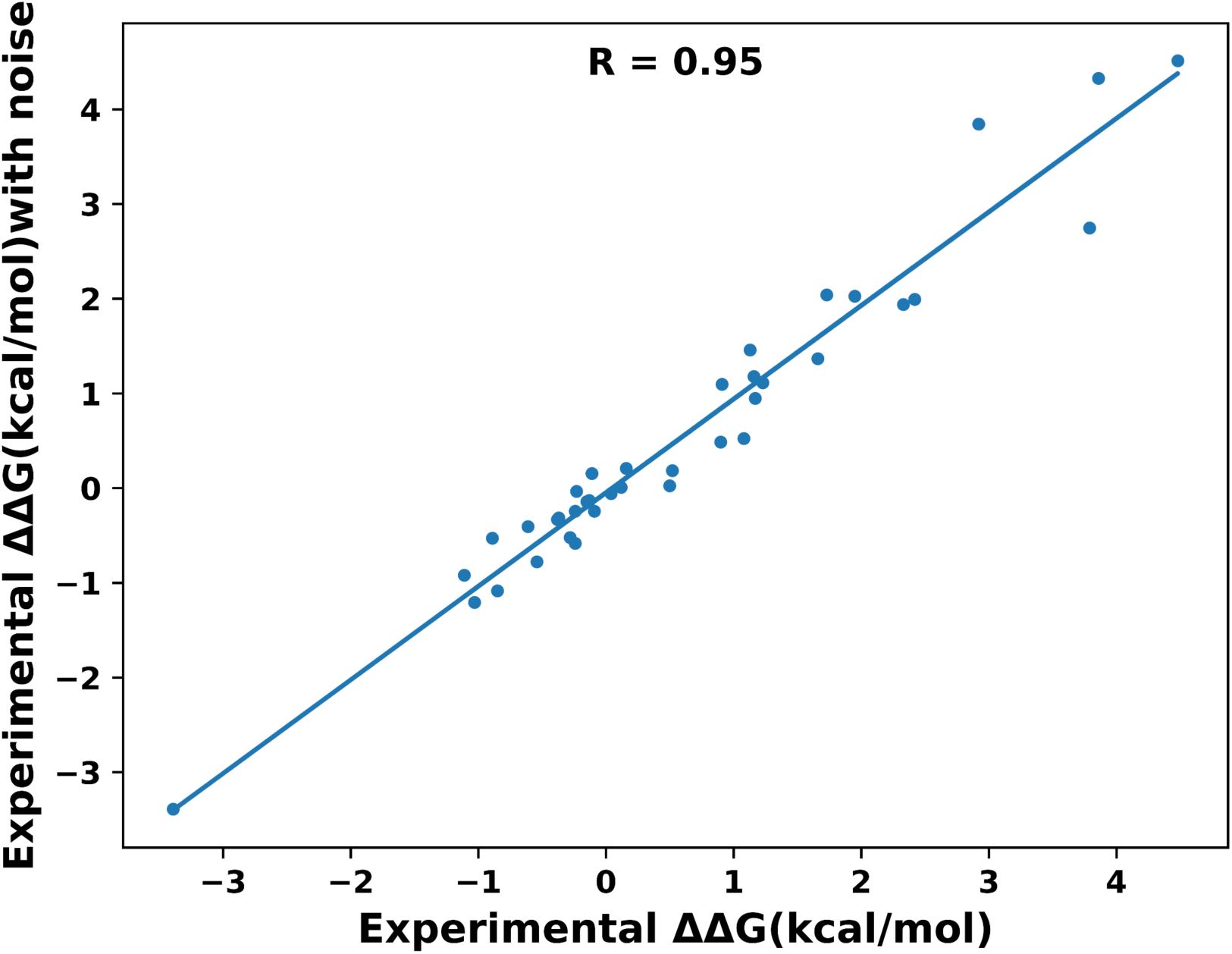
Influence of Noise on experimental data. Correlation between experimental ΔΔG_bind_ values and the same values with noise added according to the standard deviation measured for each data point. (data for colicin/DNAse complexes (PBD ID 2WPT and 1EMV).

**Supplementary Figure 3:**
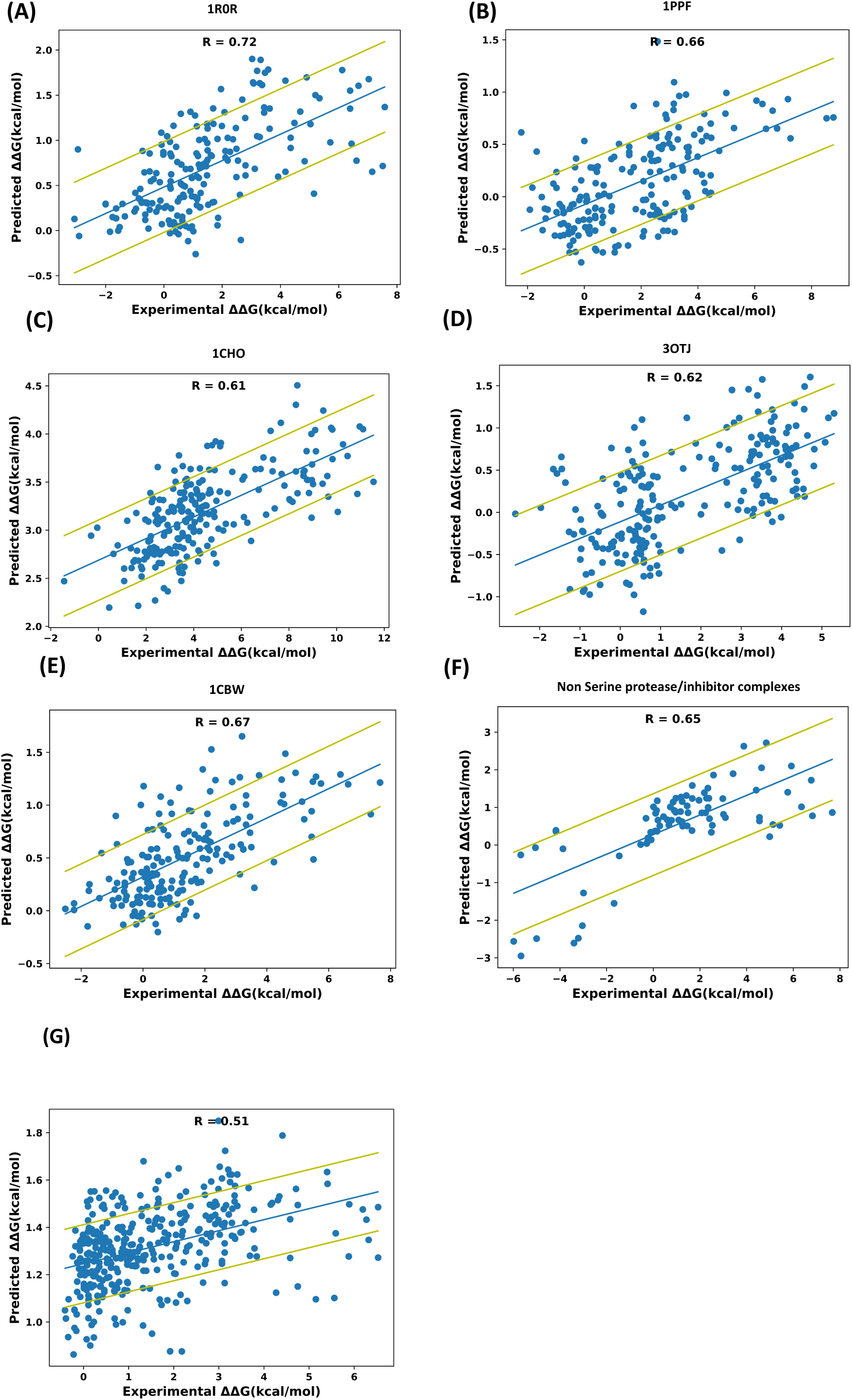
Predicting ΔΔG_bind_ for single mutations. Correlation between experimental and predicted ΔΔG_bind_ values after the model was trained on the whole dataset excluding the data for the PDB file under evaluation (A) a complex between Ovomucoid and subtilisin carlsberg (PDB ID 1R0R), (B) a complex between Ovomucoid and Human Leukocyte elastase (PDB ID 1PPF), (C) a complex between Ovomucoid and Alpha-Chymotrypsin(PDB 1CHO), (D) a complex between BPTI and Trypsin (PDB ID 3OTJ), (E) a complex between BPTI and Chymotrypsin (PDB ID 1CBW). (F) Non serine protease/inhibitor complexes (PDB IDs : 1CSE, 1CT2, 1EMV, 1S1Q, 1SBB, 1SGD, 1CT2). The blue line represents the best linear fit of the data. (G) A complex between Angiotensin-converting enzyme 2 (ACE2) and Spike protein S1 (PDB ID 6M0J). The yellow lines represent one standard deviation above and below the fitted line.

**Supplementary Figure 4:**
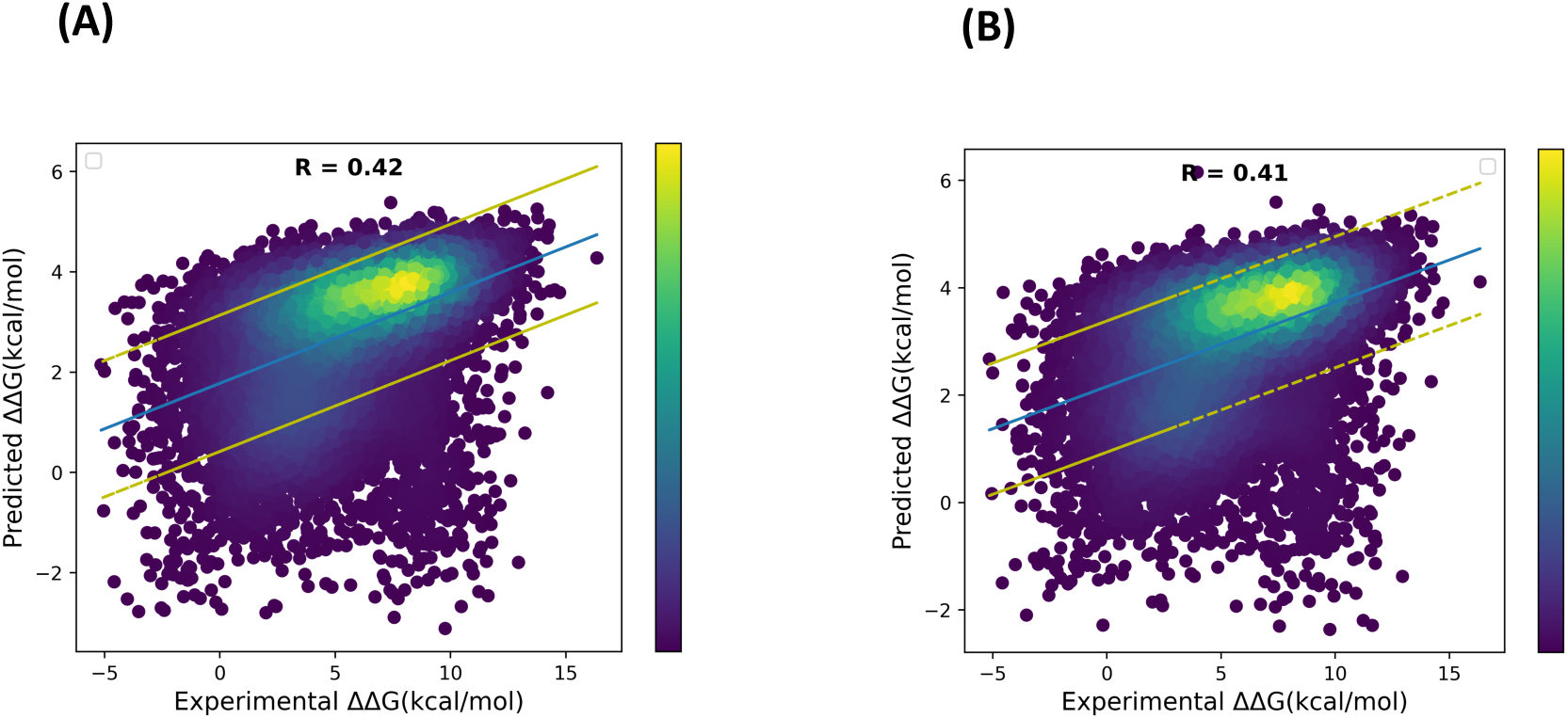
Predicting ΔΔG_bind_ for double mutations (A) Correlation between experimental and predicted ΔΔG_bind_ when model was trained on double mutations belonging to the BPTI /Chymotrypsin complex (PDB ID 1CBW) and tested on double mutations belonging to the BPTI/bovine Trypsin complex (PDB ID 3OTJ). (B) Correlation between experimental and predicted ΔΔG_bind_ when model was trained on the whole dataset of double mutants and tested on double mutations belonging to the BPTI/ bovine Trypsin complex (PDB ID 3OTJ). The blue line represents the best linear fit of the data. The yellow lines represent one standard deviation above and below the fitted line. The points are colored according to their local density, with the color bar indicating the density scale. Higher density areas (yellow color) represent regions where data points are more concentrated."

**Supplementary Figure 5:**
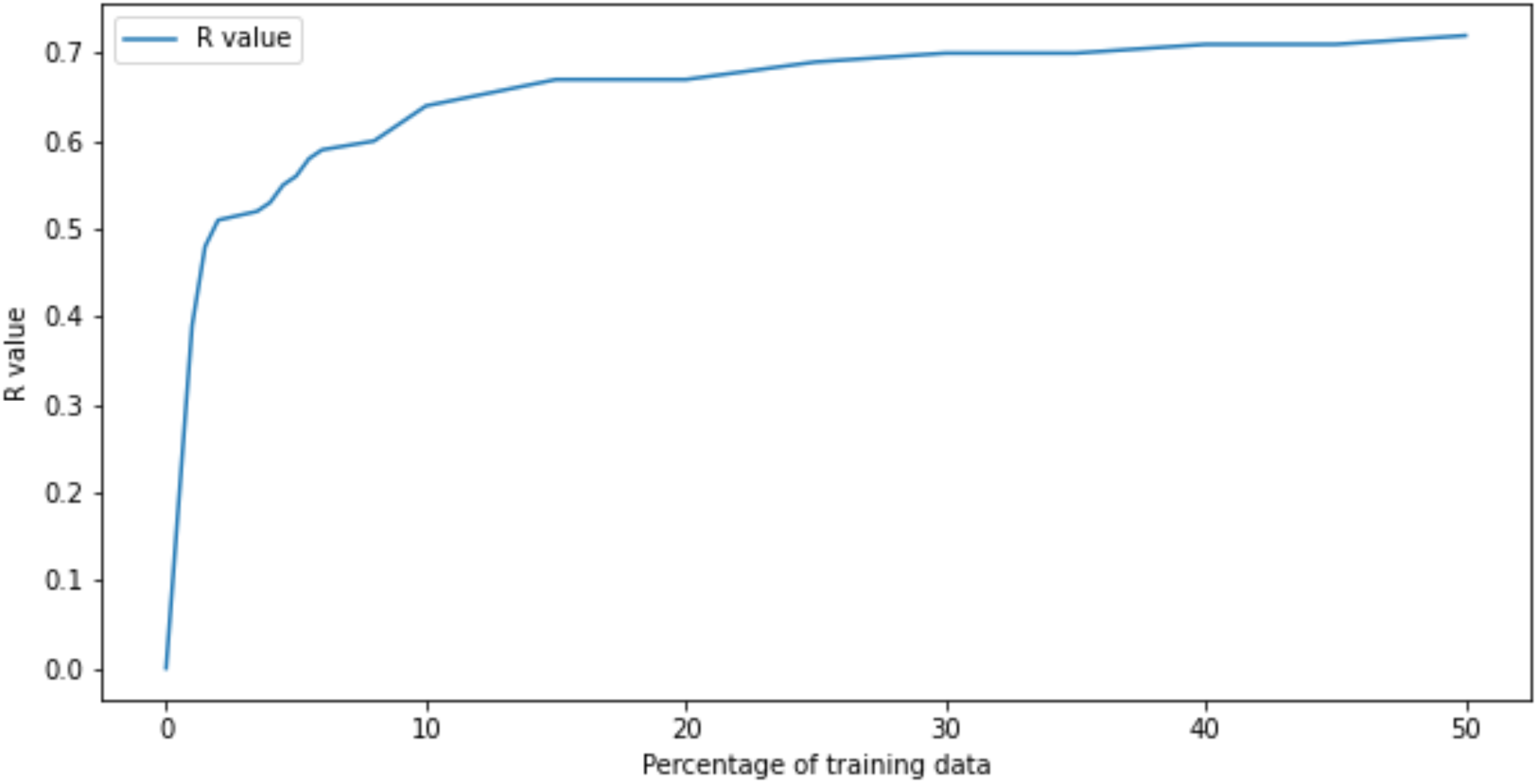
Training data vs Correlation. Graph illustrates the impact of increasing the percentage of training data on the R value between experimental and predicted ΔΔG_bind_.

**Supplementary Figure 6.**
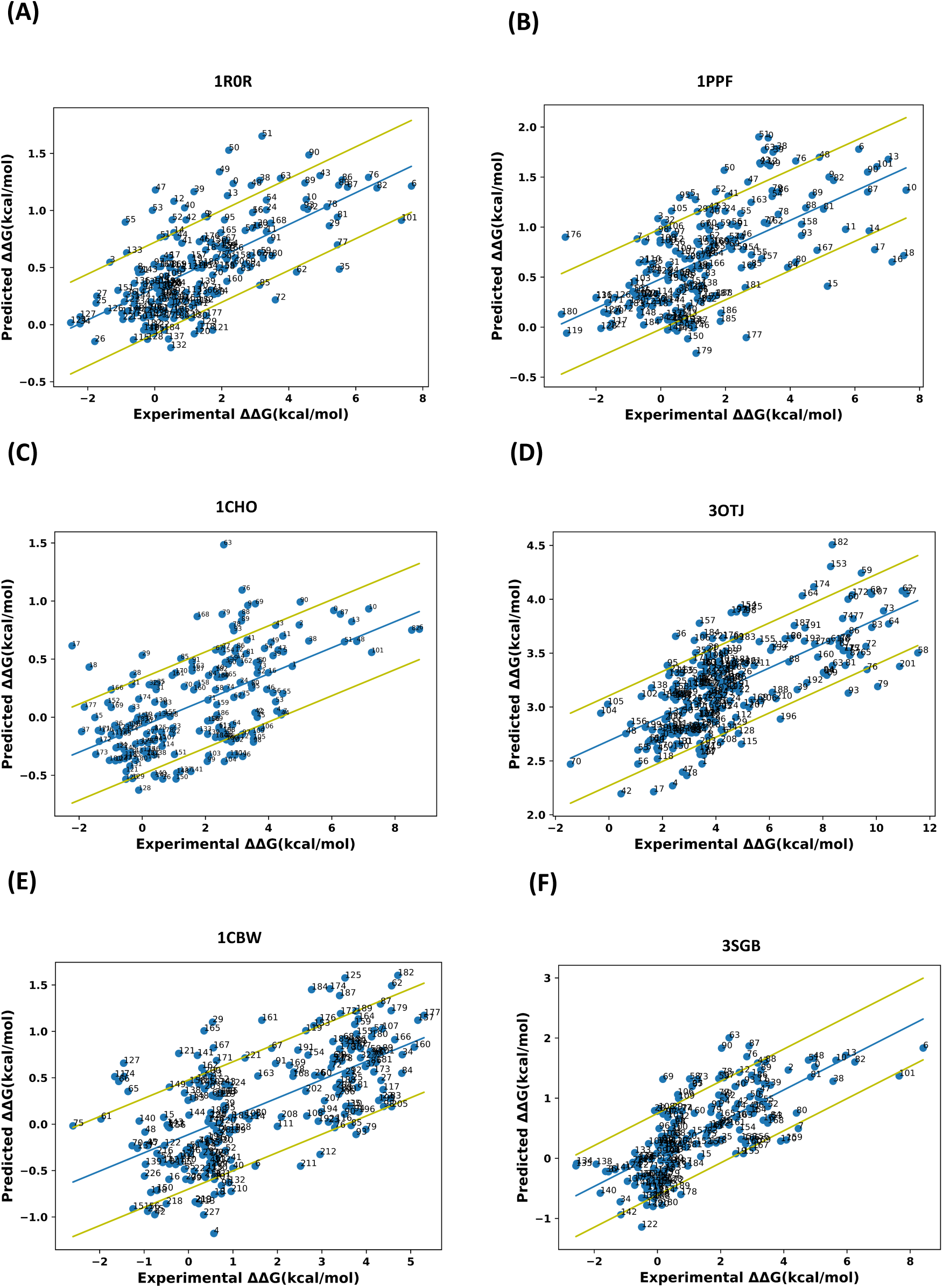
Analysis of the outliers for the six PDBs. (A) PDB ID 1R0R, (B) PDB ID 1PPF, (C) PDB 1CHO), (D) Mutation positions in the complex between BPTI and Trypsin (PDB ID 3OTJ. (E) PDB ID 1CBW, (F) PDB ID 3SGB. ProBASS was trained on the whole dataset excluding the test PDB file and predictions were made. The blue line represents the best liner fit to the data and the yellow lines correspond to one standard deviations from the fitted line. The mutations lying above and below the one-standard-deviation line were numbered and analyzed in the context of the structure. See Supplementary data for mutation description, where mutations predicted to be overly disruptive to the complex are colored in red and mutations predicted overly stabilizing for the PPI are colored in cyan. Outliers. Xlxm file is available in the ProBASS repository: (https://github.com/sagagugit/ProBASS).

